# A mathematical model for the efficient control of the New World screwworm

**DOI:** 10.64898/2026.06.21.733615

**Authors:** Rosalio Reyes, Rafael A. Barrio

## Abstract

An outbreak of New World screwworm has recently been spreading across Mexico, after more than 30 years of absence. The sterile insect technique, which consists of the massive release of sterilized males, has proven to be one of the most efficient methods for controlling the screwworm pest. However, given the limited number of sterile males available, improving the release strategy is critical. We propose a mathematical model of population dynamics adapted to the biology of *Cochliomyia hominivorax* and derive a feedback control function to determine the number of sterile males to release. We further construct a Luenberger observer to estimate wild fly populations from infected animal counts—the variable monitored by Mexican sanitary authorities—enabling field implementation of the control function. We show that eradication is achievable within approximately 60–100 weeks and that eradication time is governed primarily by the intrinsic biology of the system rather than by infestation magnitude. We then extend the model to a spatially explicit framework and show that when sterile male releases are applied at the outbreak focus and within a 120 km radius, eradication of the pest is attainable.

## Introduction

In November 2024, the first case of New World screwworm (NWS) was detected in the state of Chiapas, Mexico, on the border with Guatemala [1], after 30 years of being NWS-free [1]. To date, more than 11,000 cases have been reported [1], and the pest has reached the state of Tamaulipas [2], on the border with the United States. Therefore, it is a matter of urgent need to take actions to control this pest before it spreads further across the continent, but the resources and techniques available are limited.

The main NWS species affecting the American continent is *Cochliomyia hominivorax* [3]. It primarily infects mammals, including humans, but it can also infect birds [3]. Within the current outbreak in Mexico, infections by *Lucilia sericata, Cochliomyia macellaria*, and *Lucilia* sp. have also been reported [1]. *Cochliomyia hominivorax* is the primary infector, responsible for the initial infection of animal wounds. The other species only infect wounds after the primary infection has occurred [4]; therefore, the control of *C. hominivorax* is a prerequisite for overall pest control.

The sterile insect technique (SIT) is a biological control method that aims to reduce the birth rate through the release of sterile males [5, 6]. In NWS, this technique is particularly efficient because females in this species mate only once [7]. In fact, the first large-scale successful field eradication using the SIT targeted the New World screwworm on the island of Curaçao [8].

Previous theoretical works have proposed mathematical models of population dynamics describing the SIT [9–11]. Recent studies focused on mosquito pests that transmit dengue, malaria and Zika have used control theory to show that the SIT induces global stability [10, 11]. However, these models do not determine suitable sterile male release strategies for pest eradication under limited resources. Moreover, spatially explicit models incorporating feedback-based or resource-aware control strategies for NWS remain scarce, despite the need to account for the geographic spread of the pest.

In the present work, we propose a mathematical model to address this problem. The work is organized as follows. In the first section we propose a local population dynamics model for the New World screwworm. In the second section we determine the feedback control function for the release of sterile insect males. In the third section, we propose a Luenberger observer that allows the total wild adult fly population to be estimated from infected animal counts—a record maintained by Mexico’s National Service for Agro-Alimentary Health, Safety and Quality (Servicio Nacional de Sanidad, Inocuidad y Calidad Agroalimentaria, SENASICA)—enabling the implementation of the control function in the field. Finally, in the fourth section, we extend the model to a spatially explicit framework and show that the same control function is capable of driving the pest to extinction even under conditions of dispersal between neighboring regions. We apply the model to the specific case of the Mexican state of Chiapas.

## Methods

### Local mathematical model

Based on biological data, we devised a mathematical model to describe the local population dynamics of the organism. *Cochliomyia hominivorax* is a holometabolous insect, meaning it undergoes complete metamorphosis consisting of four stages: egg, larva, pupa, and adult. The larval stage is an obligate parasite that requires a living host to complete its development [7, 12].

Due to the characteristics of the life cycle described above and the effect on population spread, we consider only the adult stage of this pest. Therefore, we describe the dynamics through the following system of differential equations:

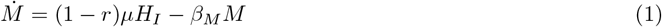

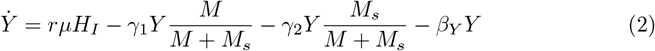

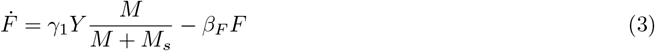

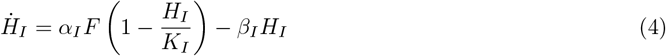

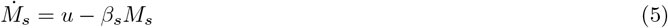

where *M, Y, F, H*_*I*_, and *M*_*s*_ represent the populations of fertile males, virgin females, mated females, infected hosts, and released sterile males, respectively. The parameters used to simulate the equations are listed in Table 1 and derived in the section Model parameter estimation. *µ* is the adult fly birth rate; *β*_*M*_, *β*_*s*_, *β*_*Y*_, and *β*_*F*_ are the death rates of fertile males, sterile males, virgin females, and mated females, respectively; *β*_*I*_ is the recovery rate of infected hosts; *r* is the proportion of females in the offspring; *γ*_1_ and *γ*_2_ represent the mating effectiveness with fertile and sterile males, respectively, where we set *γ*_2_ = *γ*_1_*/*2; and *K*_*I*_ is the carrying capacity of infected hosts. Numerical integration of the local model was performed in Python using the`solve_ivp` function from the `scipy.integrate` package, with the RK45 method and adaptive step size.

**Table 1.**
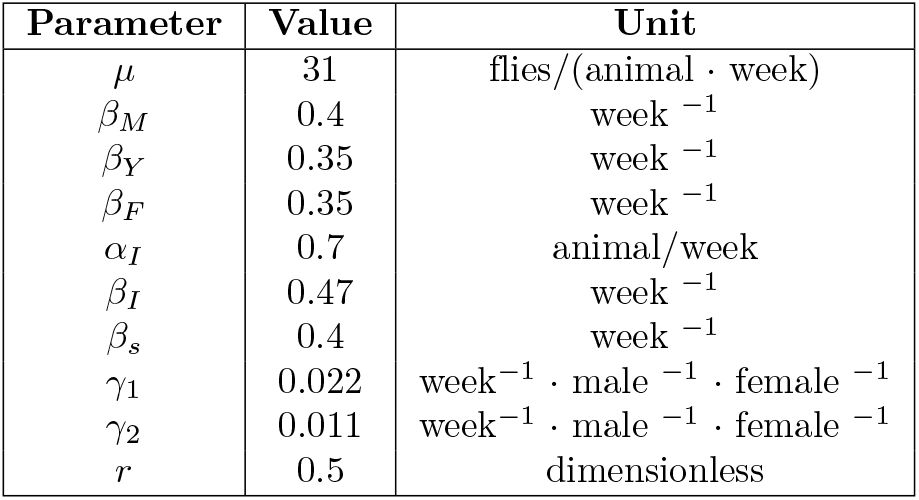
Model parameters. Parameters were estimated from the biological cycle data reported in literature and information published by SENASICA in the Avances 01–10 [1, 3]. In the Unit column, ‘animal’ refers to infected host. See section Model parameter estimation for details.

| Parameter | Value | Unit |
| --- | --- | --- |
| $\mu$ | 31 | flies/(animal · week) |
| $\beta_M$ | 0.4 | week<sup>−1</sup> |
| $\beta_Y$ | 0.35 | week<sup>−1</sup> |
| $\beta_F$ | 0.35 | week<sup>−1</sup> |
| $\alpha_I$ | 0.7 | animal/week |
| $\beta_I$ | 0.47 | week<sup>−1</sup> |
| $\beta_s$ | 0.4 | week<sup>−1</sup> |
| $\gamma_1$ | 0.022 | week<sup>−1</sup> · male<sup>−1</sup> · female<sup>−1</sup> |
| $\gamma_2$ | 0.011 | week<sup>−1</sup> · male<sup>−1</sup> · female<sup>−1</sup> |
| $r$ | 0.5 | dimensionless |

### Model parameter estimation

This section is divided into three subsections: biological facts regarding NWS, assumptions derived from these biological observations, and the resulting parameter values used in the model. A condensed summary of the information is provided in Table 1.

#### Biological facts

- A female hatch lays between 200 and 400 eggs per clutch [12].
- A female lays between two and three egg batches throughout her lifetime [12].
- Two cases of myiasis caused by NWS were reported. In one animal, 21 larvae were counted, whereas in another case, 50 larvae were reported [1, 13].
- In *Drosophila melanogaster*, the survival rate from larva to adult ranges between 0.74 and 0.97 at 25^*◦*^C [14].
- The infection process of NWS lasts between 15 and 17 days [12].
- The egg hatching time is approximately 12 hours [3].
- The larval stage lasts between 5 and 6 days [3].
- The pupal stage lasts between 7 and 10 days [3].
- The lifespan of males ranges between 14 and 21 days [12].
- The lifespan of females ranges between 10 and 30 days [12].
- A male mates approximately 5 to 6 times during its lifetime [12].
- The mating fitness of sterile males (*γ*_2_) is approximately half that of wild males (*γ*_1_) personal communication from SENASICA personnel.

#### Model assumptions

- We assume that a female lays 300 eggs per clutch and oviposits three times during its lifetime.
- Since a female lays eggs on approximately three occasions during its lifetime, we assume that each female infects an average of 2.5 animals.
- The reported NWS infection cases are assumed to originate from a single female fly in each case. Under this assumption, since a female deposits approximately 480
- eggs per animal, only 20% of the eggs survive to the larval stage, corresponding to approximately 96 larvae.
- The infection delay is assumed to be 2.1 weeks.
- Based on the *Drosophila* case, we assume a larva-to-adult survival rate of 0.7.
- The egg hatching time is assumed to be 0.07 weeks.
- The larval stage duration is assumed to be 0.86 weeks.
- The pupal stage duration is assumed to be 1.21 weeks.
- The lifespan is assumed to be the same for virgin and mated females, equal to 2.86 weeks.
- The lifespan is assumed to be the same for wild and sterile males, equal to 2.5 weeks.
- We assume a 1:1 sex ratio at birth.

#### Derived parameters

- A female lays approximately

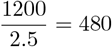

eggs per animal.
- From the 480 eggs deposited on an animal, we estimate

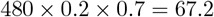

adult flies.
- The parameter

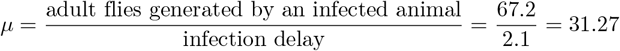

flies/(animal*·*week).
- The infection rate is given by

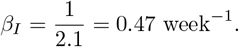
- The mortality rates of wild and sterile males are

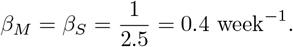
- The mortality rates of virgin and mated females are

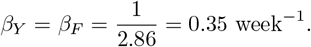
- The infection rate per female is estimated as

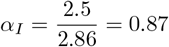

animals/week.
- The mating rate of males is estimated as

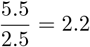

matings/(week*·*male).
- The mating parameter is estimated as

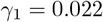

week^*−*1^male^*−*1^female^*−*1^.
- Since sterile males are assumed to have half the mating fitness of wild males,

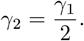
- *r* = 0.5

### How many sterile males should be released?

Let *x* = (*M, Y, F, H*_*I*_, *M*_*s*_)^*T*^. We write the model Eq (1)–Eq (5) in the form

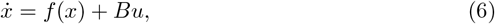

where *f* (*x*) is the vector field whose first four components correspond to Eq (1)–Eq (4), the fifth component is *−β*_*s*_*M*_*s*_, and *B* = (0, 0, 0, 0, 1)^*T*^.

A system is said to be controllable at time *t*_0_ if it is possible, by means of an unconstrained control vector, to transfer the system from any initial state *x*(*t*_0_) to any other state in a finite time interval [15]. For a linear system of the form 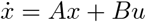, where *x∈* ℝ^*n*^ is the state vector, *A ∈* ℝ^*n×n*^, and *B∈* ℝ^*n×m*^, the integer *n* denotes the number of state variables and *m* the number of control inputs. The system is controllable if the controllability matrix [*B* |*AB* |*· · ·* |*A*^*n−*1^*B*] has full rank [15]. We evaluated the controllability of the linearization of model Eq (6) at 50,000 points in the state space, with each component of *x* and the carrying capacity varying in the range (0, 10^8^), obtaining all sampled points yielded a full-rank controllability matrix. Since controllability of the linearized system implies local controllability of the nonlinear system [16], these results suggests that the model is locally controllable over a broad region of the state space and that suitable control function for SIT may exist. At each point, the Jacobian *A* = *∂f/∂x* was computed symbolically and evaluated numerically; the rank of was then assessed using the ctrb function of the MATLAB Control System Toolbox.

We design a control function *u* (see Eq. (6)), defined as the sterile male release ratio, to determine the number of sterile males required for pest eradication. We define the extinction time *t*_*E*_ as the instant at which, after the introduction of sterile males, all populations (*M, Y, F, H*_*I*_) fall below 1. The total number of sterile males released is defined as 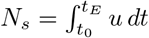. We evaluated the tradeoff between reducing *N* and shortening *t*_*E*_ by numerically exploring control functions of the form *u* = *Gx*, where *G ∈* ℝ^1*×*5^ is usually known as a gain matrix, in this case it is a vector of coefficients that weights the state variables to compute the control action.

### Measurement of control variables in the field

In practice, counting the number of wild males is difficult, primarily because sterile NWS males are not marked to distinguish them from fertile ones. Moreover, it is difficult to determine whether a captured female is virgin or mated. This hinders the direct implementation of the proposed control functions, which depend on *M* and *Y*. However, it is possible to estimate the wild populations from an observable variable.

A system is said to be completely observable if every state *x*(*t*_0_) can be determined from the observation of the output *y*(*t*) over a finite time interval *t*_0_ *≤ t ≤ t*_1_ [15]. We propose using *H*_*I*_, the number of infected animals, as the output variable to estimate the remaining populations, since SENASICA has a historical record of monitoring *H*_*I*_ [2]. To this end, we write the model in the form

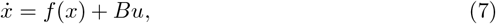

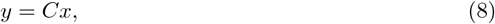

where *y* is the measured output and *C* = (0, 0, 0, 1, 0). Since *M*_*s*_ is known to the operator, only the wild populations (*M, Y, F, H*_*I*_) need to be estimated. Therefore we treat *M*_*s*_ as a known parameter and reduce the system to

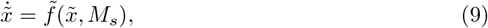

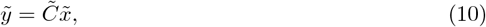

where 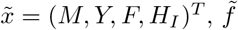 consists of Eq (1)–Eq (4), and 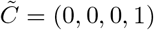.

For a linear system of the type 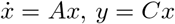, then complete observability holds if and only if the *nm × n* matrix 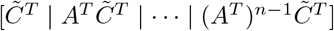 has rank *n*. We evaluated the observability of the linearized reduced subsystem at 10,000 points in the state space, with each component varying in the range (0.1, 10^8^) and *K*_*I*_ = 10^8^. Full rank was obtained in all cases. These results suggest that the wild population variables may be locally reconstructible from measurements of *H*_*I*_ over a broad region of the state space [15], indicating that all four wild populations are observable from the measurement of *H*_*I*_.

### Numerical implementation of the observer

To estimate the wild populations and use these estimates in the control function, we construct an extended Luenberger observer [16]. The coupled system consists of the real model and the observer, solved simultaneously:

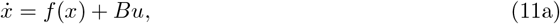

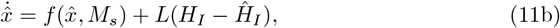

where *x* = (*M, Y, F, H*_*I*_, *M*_*s*_)^*T*^ is the real state, 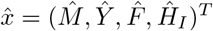 is the estimated state, and *L ∈* ℝ^4*×*1^ is the observer gain. The term *L*(*H*_*I*_ *−Ĥ*_*I*_) corrects the estimate using the difference between the real and estimated measurements. The gain *L* is recomputed at each time step by local linearization around the current observer state vector 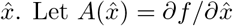 denote the Jacobian matrix evaluated along the observer trajectory. Then, the dynamic gain *L* is obtained as

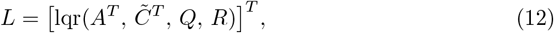

with *Q* = 0.01 *I*_4*×*4_, where *I*_4*×*4_ denotes the 4 *×*4 identity matrix, and *R* = 1. Here, `lqr` is the MATLAB Control System Toolbox function that computes the optimal state-feedback gain for a continuous-time linear quadratic regulator problem by solving the associated algebraic Riccati equation [15]. Under standard observability assumptions, this choice of *L* yields asymptotically stable estimation error dynamics, i.e.,

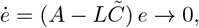

where the estimation error vector is defined directly as 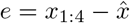, with *x*_1:4_ = (*M, Y, F, H*_*I*_)^*T*^ representing the sub-vector of the real wild populations.

The coupled system Eq (11) was integrated using a fixed step-size Euler scheme with Δ*t* = 0.1 weeks, to avoid artifacts.

### Spatial model

We extend model Eq (1)–Eq (5) by defining a particular geographical region with a square grid. In each cell we define a local model and propagation between neighbouring cells is described by adding a diffusion term to each of the variables *M, Y, F*, and *M*_*s*_, which incorporate the spatial spread of adult NWS flies between neighbouring cells:

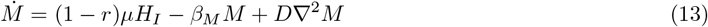

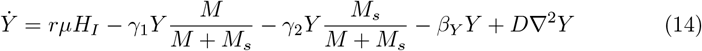

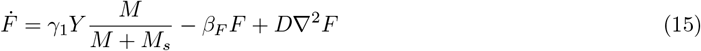

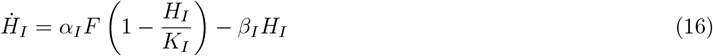

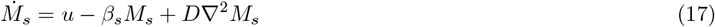

where *D* is the diffusion coefficient of adult flies. For simplicity, the same diffusion coefficient was assumed for males, females, and sterile males. The Laplacian was discretized using a standard five-point finite-difference stencil. We investigate the case of the Mexican state of Chiapas. A 50 *×*50 grid was used, where each cell represents approximately 8 *×*8 km. This spatial resolution was selected for computational efficiency and does not correspond to any particular biological or geographical scale. Dirichlet boundary conditions (*M* = *Y* = *F* = *M*_*s*_ = 0 on *∂*Ω) are imposed to model the absence of flies beyond the domain. These boundary conditions are sufficient because long-range re-introductions from outside the domain are incorporated separately through stochastic seeding events.

In order to define the diffusion coefficient, we initially set *D* = 0.2 (in grid-cell units per week), corresponding to a physical value of

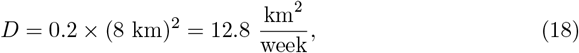

which implies a root-mean-square dispersal distance of [17]

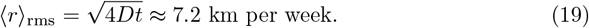

This value is consistent with empirical mark-recapture data for *Cochliomyia hominivorax* reported by [18]. Under favorable environmental conditions—warm temperatures and high host density, characteristic of the livestock-producing lowlands of Chiapas—the 90th-percentile dispersal radius was estimated at 2.85 km for females and 1.25 km for males per observation period. Under less favorable conditions, these values increase up to 22.07 km and 9.64 km, respectively. The chosen diffusion coefficient places the effective weekly dispersal distance (7.2 km) between these two regimes, representing an intermediate scenario appropriate for the heterogeneous landscape of Chiapas, where both dense agricultural areas and less suitable terrain coexist. However, rather than fixing the diffusion coefficient at this value, we further refine the analysis by exploring D over the interval (0, 0.3). Simultaneously, we investigate the effect of a stochastic pest-propagation term. The procedure used to estimate these quantities is described below.

Time integration was performed with a forward Euler scheme (Δ*t* = 0.1 weeks) implemented in Python using vectorized NumPy array operations.

The carrying capacity *K*_*I*_ at each grid cell was estimated from livestock density data from the 2022 Mexican Agricultural Census by the National Institute of Statistics and Geography (Instituto Nacional de Estadística y Geografía, INEGI) [19]. Cells with negligible livestock density (*K*_*I*_ *<* 10^*−*6^) were treated as inactive and their derivatives set to zero in order to avoid numerical issues.

Independent re-outbreaks in distant cells were modeled as stochastic events. At each time step Δ*t*, a new infestation focus was seeded at a random cell with a probability *p* = *λ*_seed_Δ*t*, where *λ*_seed_ represents the mean seeding rate. This Monte Carlo approach enables the model to capture sporadic re-introductions from outside the controlled zone as well as stochastic pest establishment in distant areas.

To calibrate the model, a bi-dimensional grid search was conducted across 2,500 uniformly spaced parameter combinations by partitioning the diffusion coefficient interval *D ∈* [0.002, 0.300] and the seeding rate interval *λ*_seed_ *∈* [0.05, 0.50] into 50 levels each. The simulation horizon spanned 59 weeks, synchronized with weekly field records from the SENASICA database covering the period from November 20, 2024, to January 7, 2026. To account for stochastic variability, 30 independent replicates were executed for each parameter pair (*D, λ*_seed_).

The spatio-temporal fit was evaluated using the spatial Jaccard index (*J*) as the primary objective function, defined at each empirical checkpoint *t* as:

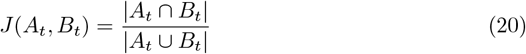

where *A*_*t*_ represents the set of empirically infected municipalities and *B*_*t*_ denotes the set of dynamically simulated infected municipalities, both defined by an active host threshold of *H*_*I*_ *≥*1. The parameter configuration that maximized the mean Jaccard index across all temporal field checkpoints, driving the score closest to 1, was selected as the optimal fit.

The calibration procedure identified an optimal parameter configuration of *D* = 0.002 and *λ*_seed_ = 0.1143, yielding a mean Jaccard index of 0.1746. Using these optimized parameters, the model’s overall performance was assessed through cross-validation by computing the Mean Absolute Error (MAE) over 50 independent runs for both the total number of affected municipalities and the overall fraction of infected state area. Under this optimal parameter regime, the model exhibited good predictive performance, with a mean error of 6.1 municipalities (standard deviation = 1.69) and a mean absolute error of 12.24% in the infected-area fraction (standard deviation = 0.0153).

## Results

### Low infestation levels are sufficient to trigger uncontrolled population growth

In the uncontrolled model, i.e., when *u* = 0 and *M*_*s*_ = 0, the system Eq (1)–Eq (4) has three fixed points. The components *M* ^*\**^, *Y* ^*\**^, and *F* ^*\**^ are expressed as functions of 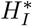:

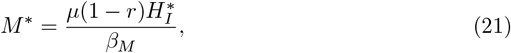

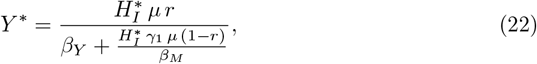

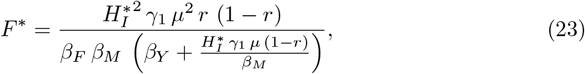

where 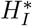 is a solution of

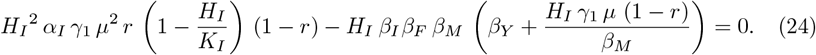

Using the parameters in Table 1 and a carrying capacity *K*_*I*_ = 1000, we obtain the following fixed points:

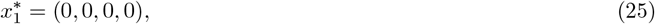

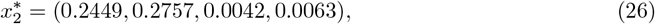

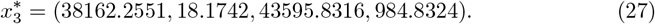

Table 2 shows the eigenvalues of the Jacobian of the uncontrolled model, demonstrating that 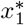 and 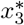 are stable fixed points, while 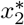 is unstable. Therefore, initial conditions in a neighborhood of 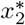 that are not contained in its stable manifold lead to uncontrolled pest growth.

**Table 2.**
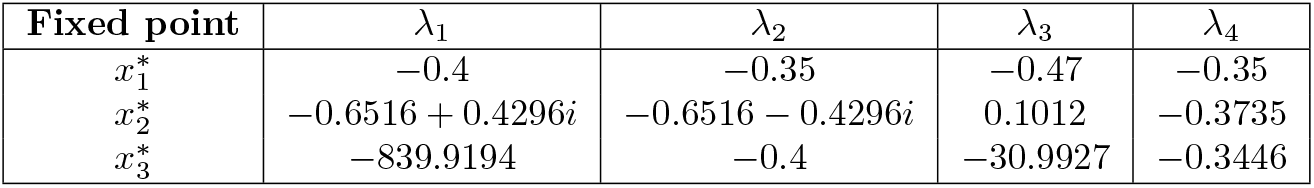
Eigenvalues associated with each fixed point.

While male individuals (*M*) and unmated females (*Y*) alone cannot initiate pest population growth from an all-zero state, mated females (*F*) and infected hosts (*H*_*I*_) possess the capacity to generate new individuals, thereby triggering pest establishment. We numerically explored the critical thresholds at which the pest either faces extinction or grows towards its carrying capacity. To achieve this, all state variables were initialized at zero except for the specific variable of interest (*F* or *H*_*I*_), and we implemented the bisection method over the domain [0, 100] with a tolerance of 10^*−*6^ to locate the exact bifurcation points. We found that the critical point for mated females is *F* ^(0)^ *≈* 0.0124 and for infected hosts is 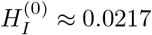, as graphically shown in Fig. 1. Observe that very low values of infection vectors are sufficient for the pest to grow indefinitely.

**Fig 1.**
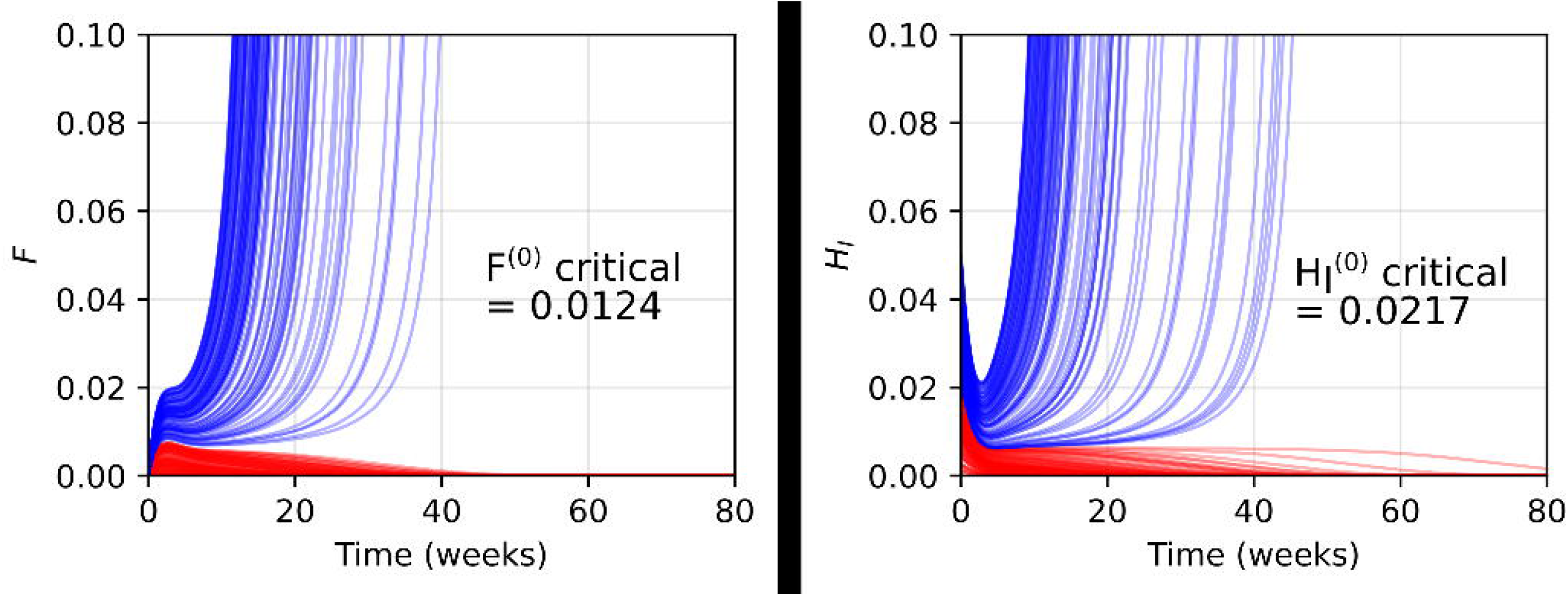
Numerical exploration of the extinction/explosion threshold. Threshold from initial conditions of *F* (a) or *H*_*I*_ (b). Blue curves correspond to solutions above the threshold (pest explosion), red curves to solutions below the threshold (pest extinction).

When we analyze the fixed points of the full controlled model, we again obtain three equilibria. The first one,

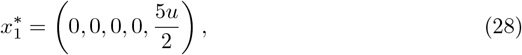

is stable. The remaining two depend on *K*_*I*_ and *u*, and take the general form:

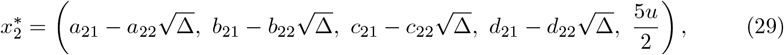

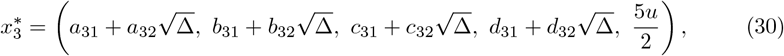

where Δ = *K*_*I*_ (160,757,041 *K*_*I*_ *−* 18,663,700 *u*) is a discriminant. *a*_*ij*_, *b*_*ij*_, *c*_*ij*_, and *d*_*ij*_ are model-derived coefficients that may depend on the parameter *K*_*I*_ but are independent of the control input *u*. Their explicit expressions are given in the Supporting information.

This expression reveals the existence of a threshold *u ≈*8.61 *K*_*I*_ at which 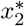 and 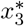 collide and annihilate in a saddle-node bifurcation, leaving 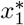 as the only equilibrium. Consequently, by choosing *u* sufficiently large, any initial state of the pest population will be naturally drawn toward the extinction equilibrium. This motivates the use of an optimal control framework to determine the conditions under which the pest is driven to extinction using a minimal number of sterile males in an optimal time.

### Feedback-based sterile male release strategies for pest eradication

To identify effective feedback control function *u* for model Eq (1)–Eq (5), we begin with a univariate analysis to determine which state variable has the greatest influence on the control. We analyze control functions of the form *G* = *n ê*_*i*_, where *n* takes integer values between 1 and 1000 and *ê*_*i*_ is a canonical basis vector of ℝ^5^, with *i ∈ {*1, 2, 3, 4*}* corresponding to the state variables *M, Y, F*, and *H*_*I*_, respectively. As shown in the first column of Fig 2, an inverse relationship between the control factor *n* and the extinction time *t*_*E*_ is observed exclusively for the entries corresponding to *M* and *Y*, where *t*_*E*_ tends asymptotically to a lower bound. In contrast, for *F* and *H*_*I*_ the trend is more gradual, requiring much larger control efforts (*n >* 500 for *F, n >* 250 for *H*_*I*_) to achieve extinction. The second column of Fig 2 reveals an inherent conflict between the objectives. Tracking the solutions shows that minimizing the extinction time *t*_*E*_ requires a higher total number of released sterile males *N*_*s*_, a trade-off that underpins the multi-objective optimization problem. Furthermore, the third column of Fig 2 demonstrates that the total number of insects used (*N*_*s*_) does not scale linearly with the gain *n*, exhibiting a high dispersion that justifies the need for an optimized bivariate combination rather than simply increasing a single parameter. From this analysis, we conclude that the entries corresponding to *M* and *Y* are the most important for control. We identified the value of *n* beyond which further increments do not produce a significant reduction in the extinction time, obtaining *n* = 99 for *M* and *n* = 56 for *Y*.

**Fig 2.**
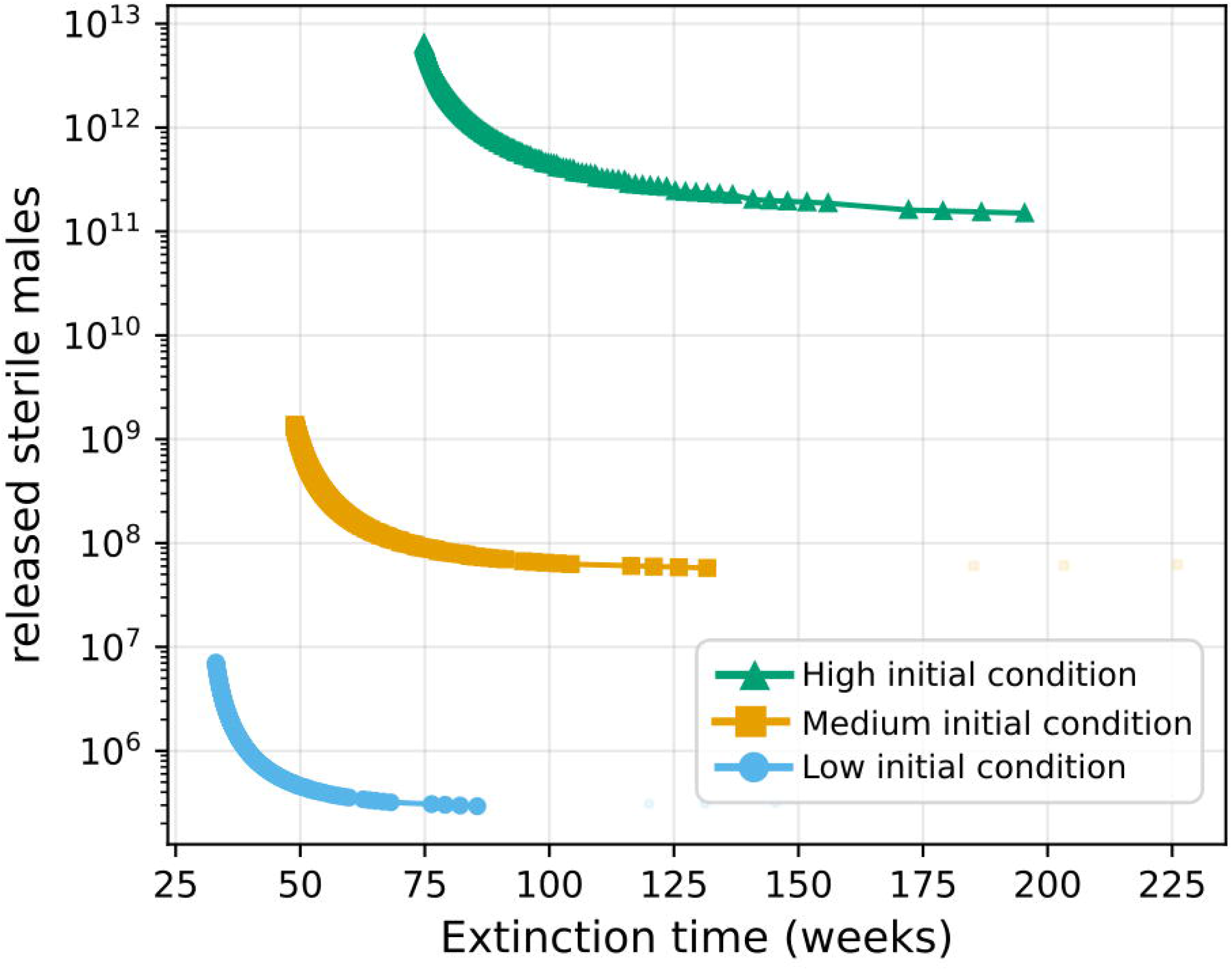
Univariate analysis of the gain matrix. Each row corresponds to a state variable (*M, Y, F, H*_*I*_). Left column: extinction time vs. factor *n*. Center column: sterile males released vs. extinction time. Right column: sterile males released vs. factor *n*.

To refine the control function, we explored the bivariate space *G* = (*n, m*, 0, 0, 0) with 0*≤ n≤* 99 and 0*≤ m≤* 56, evaluating all 100*×* 57 = 5,700 integer combinations. The exploration was carried out under three sets of initial conditions representative of different infestation scales:

- Low: (*M*_0_, *Y*_0_, *F*_0_, *H*_*I*,0_, *M*_*s*,0_) = (100, 50, 10, 500, 0),
- Medium: (*M*_0_, *Y*_0_, *F*_0_, *H*_*I*,0_, *M*_*s*,0_) = (10^4^, 5 *×* 10^3^, 10^3^, 5 *×* 10^4^, 0),
- High: (*M*_0_, *Y*_0_, *F*_0_, *H*_*I*,0_, *M*_*s*,0_) = (10^8^, 5 *×* 10^7^, 10^7^, 5 *×* 10^8^, 0).

For each combination (*n, m*) and each set of initial conditions, the system was solved numerically and *t*_*E*_ and *N*_*s*_ were recorded. Since *t*_*E*_ and *N*_*s*_ are conflicting objectives—reducing one implies increasing the other—there is no single optimal solution. Instead, the Pareto front was constructed for each set of initial conditions. A solution (*n*_1_, *m*_1_) dominates another (*n*_2_, *m*_2_) if and only if *t*_*E*_(*n*_1_, *m*_1_) *≤ t*_*E*_(*n*_2_, *m*_2_) and *N*_*s*_(*n*_1_, *m*_1_) *≤N*_*s*_(*n*_2_, *m*_2_), with at least one strict inequality. The Pareto front is the set of non-dominated solutions, representing the best possible trade-offs between both objectives.

Fig 3 shows the Pareto fronts for the three scales of initial conditions. All three curves exhibit the same structure: an inverse relationship between *t*_*E*_ and *N*_*s*_ with a hyperbolic shape. The knee of each curve is located approximately in the same extinction time range (60–100 weeks), suggesting that the minimum extinction time is determined by the intrinsic dynamics of the system rather than by the scale of the infestation. The number of sterile males, on the other hand, scales proportionally with the magnitude of the initial conditions, which is consistent with the linear structure of the control *u* = *Gx*.

**Fig 3.**
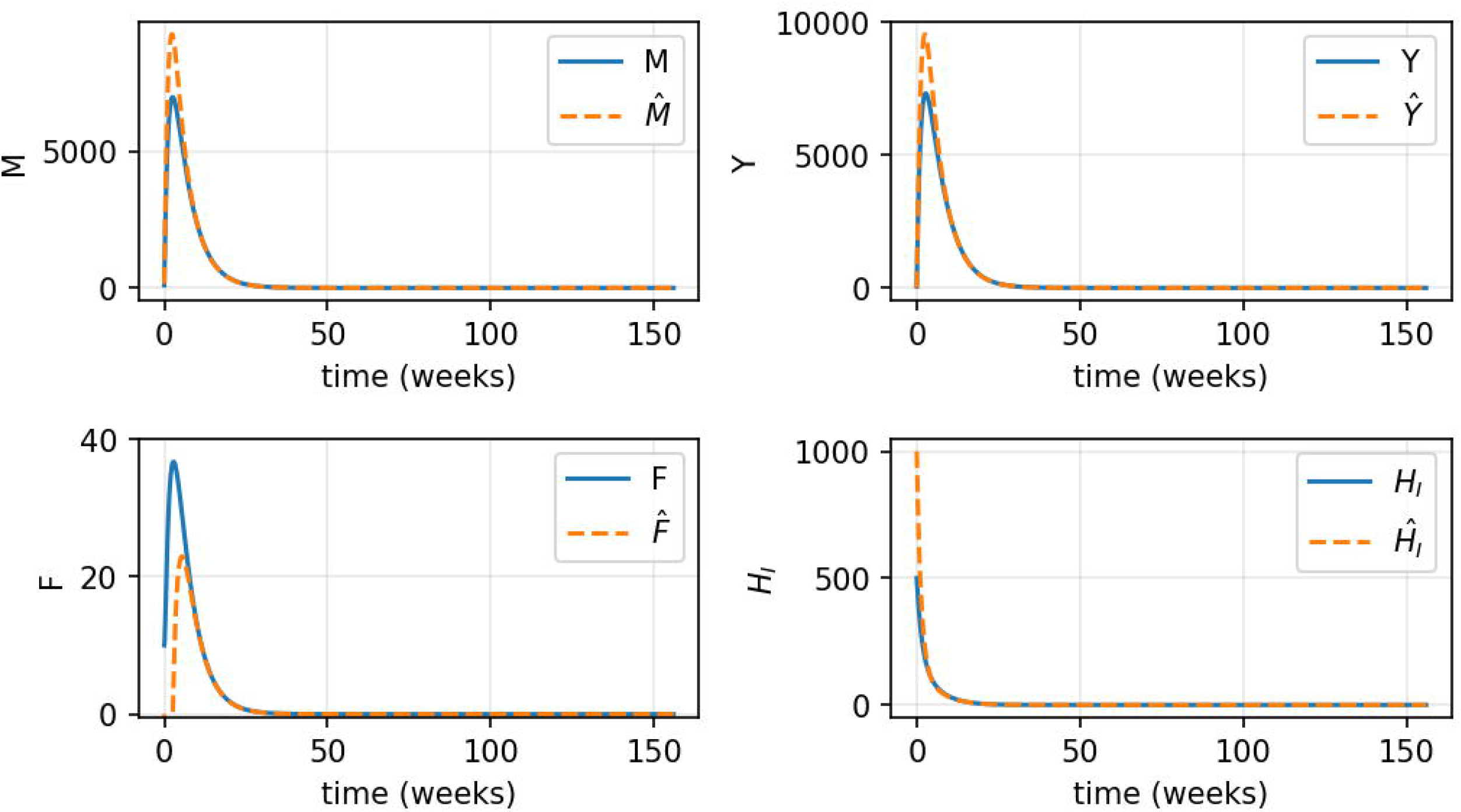
Pareto fronts for three scales of initial conditions. Low, medium, and high infestation scenarios. All three curves exhibit the same hyperbolic structure, with the knee located in the 60–100 week range.

To obtain credible predictions, we identify gain matrices capable of driving the pest population to extinction within prescribed time limits of 40, 52, 78, and 104 weeks. Among the feasible solutions satisfying each extinction-time constraint, we select the solutions with the smallest values of *N*_*s*_. As shown in Table 3, for tighter constraints such as 40 and 52 weeks, there are scenarios where, under medium or high initial infestation levels, no feasible gain matrix capable of eradicating the pest was identified.

**Table 3.**
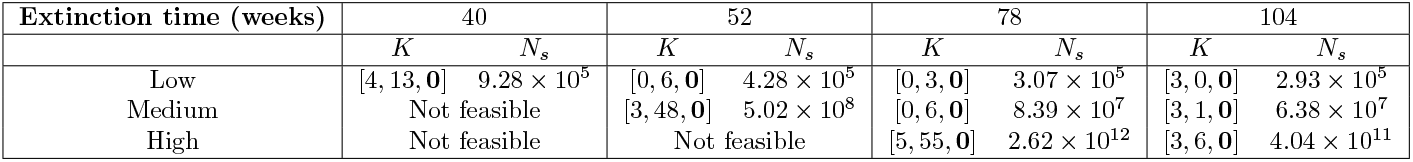
Optimal gain matrix *K* and total sterile males released (*N*_*s*_) for different maximum extinction times and initial condition scales. **0** denotes the remaining zero entries. The time constraint represents the maximum allowable extinction time. ‘Not feasible’ indicates that no gain matrix could be found to successfully drive the pest population to extinction within the constrained timeframe.

### In a simplified model, the eradication time depends on biological parameters

In order to analyze the pest extinction time, we consider a simplified model in which population growth is not limited by a carrying capacity and sterile males are introduced at a constant rate. The resulting system is

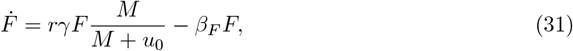

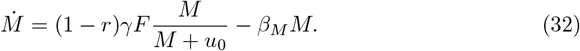

This system has two equilibrium points,

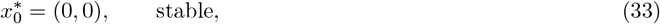

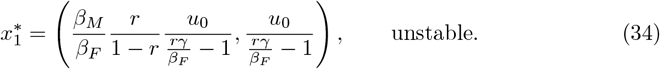

Initial conditions below 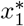 lead to extinction, whereas trajectories above 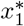 exhibit unbounded growth. The Jacobian matrix evaluated at the extinction equilibrium has eigenvectors (1, 0) and (0, 1), associated with eigenvalues −*β*_*F*_ and −*β*_*M*_, respectively. Therefore, trajectories approaching the extinction equilibrium decay at rates determined by the mortality parameters, *i*.*e*., by the biological characteristics of the pest population, consistent with our numerical predictions.

### Observer-based control successfully eradicates the pest

As discussed above, under the current NWS outbreak conditions in Mexico, it is not possible to directly measure the number of wild males or virgin females in the field. Therefore, using the observer described above, we evaluate the control function on the estimated states:

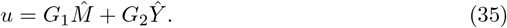

As shown in Fig 4, the control based on estimated states successfully drives the pest to extinction. This result holds for all three scales of initial conditions, provided the gain matrices reported in Table 3 are used.

**Fig 4.**
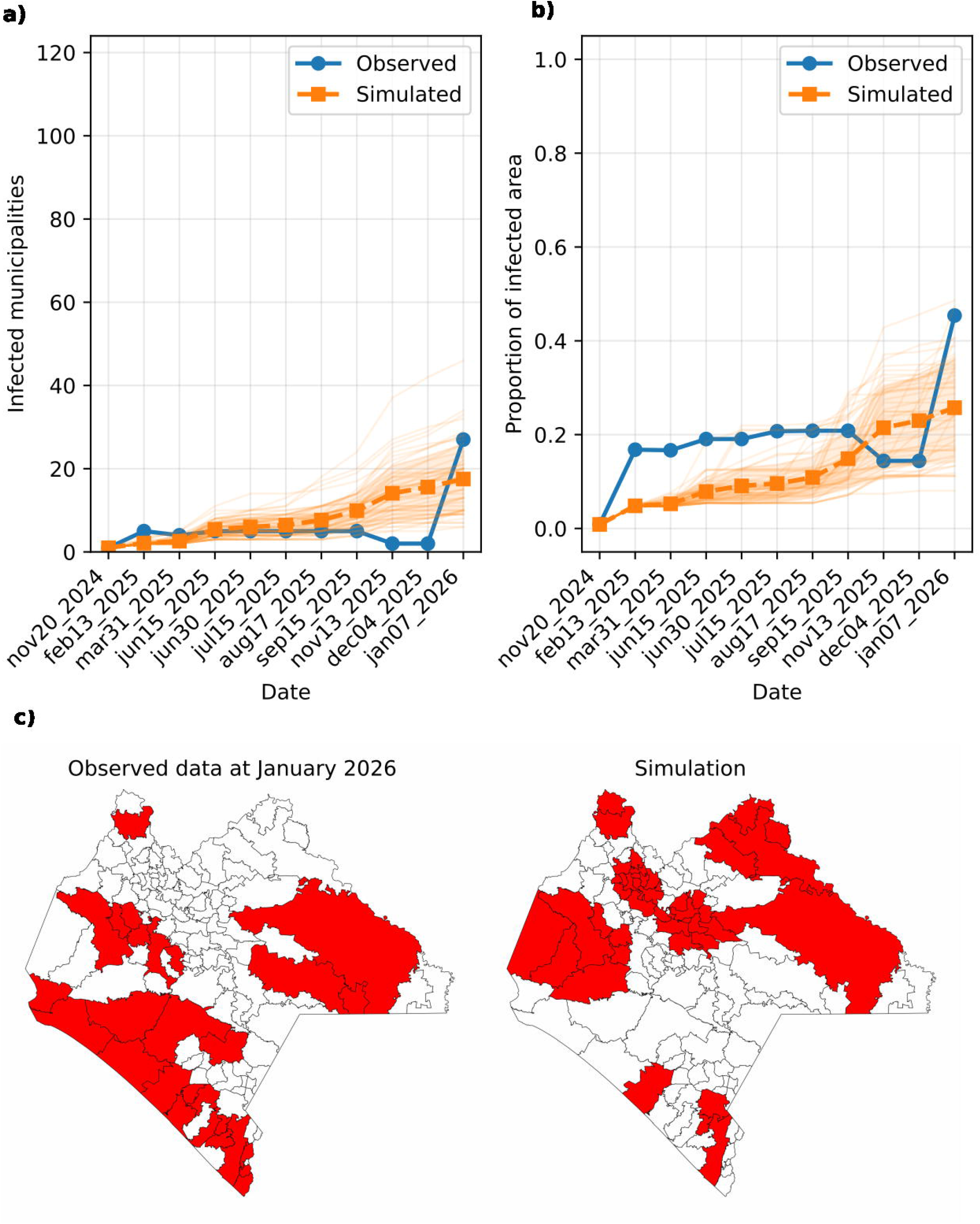
Solutions of the coupled system (real model + observer). Low initial conditions with *G* = [0, 6, **0**]. The pest is eradicated at *t*_*E*_ *≈*50 weeks, consistent with the value reported in Table 3.

### Spatial model: Application to the case of Chiapas

We implement the spatial model described above in the state of Chiapas, located in southern Mexico, on the border with Guatemala (Fig 6a).

According to SENASICA records, the initial New World Screwworm (NWS) outbreak in Chiapas was reported in the northern municipality of Catazajá [1]. Based on this epidemiological milestone, our simulation initialized the infestation front within this specific geographic focus. Concurrently, independent outbreaks across the state were dynamically incorporated via Monte Carlo stochastic seeding. The resulting spatio-temporal alignment between the empirical SENASICA data and our calibrated model, under the optimal parameters determined by the Jaccard index, is illustrated in Fig. 5. Furthermore, Fig. 5 (a)-(b) displays the simulated spatial patterns of infected municipalities and proportion of infected area. While replicating the exact stochastic configuration of field outbreaks is inherently impossible due to the probabilistic nature of the seeding process, the model captures the global macroeconomic and regional dynamics with high fidelity. As detailed in the Methods section, the framework’s performance was cross-validated using the Mean Absolute Error (MAE) for both the total count of affected municipalities and the cumulative fraction of infected state area. Under this optimal parametric regime, the model achieved a high degree of predictive accuracy, yielding a mean error of 6 municipalities and a 12% deviation in the total fraction of infected area. The simulation dynamics can be viewed in Supplementary Video S1. Although the model operates on a 50*×* 50 grid covering Chiapas, the video displays results at the municipal level. This visualization was generated by intersecting the grid with municipal borders and aggregating the total cases of all pixels within each municipality. Municipalities with at least one infected host case are highlighted in red.

**Fig 5.**
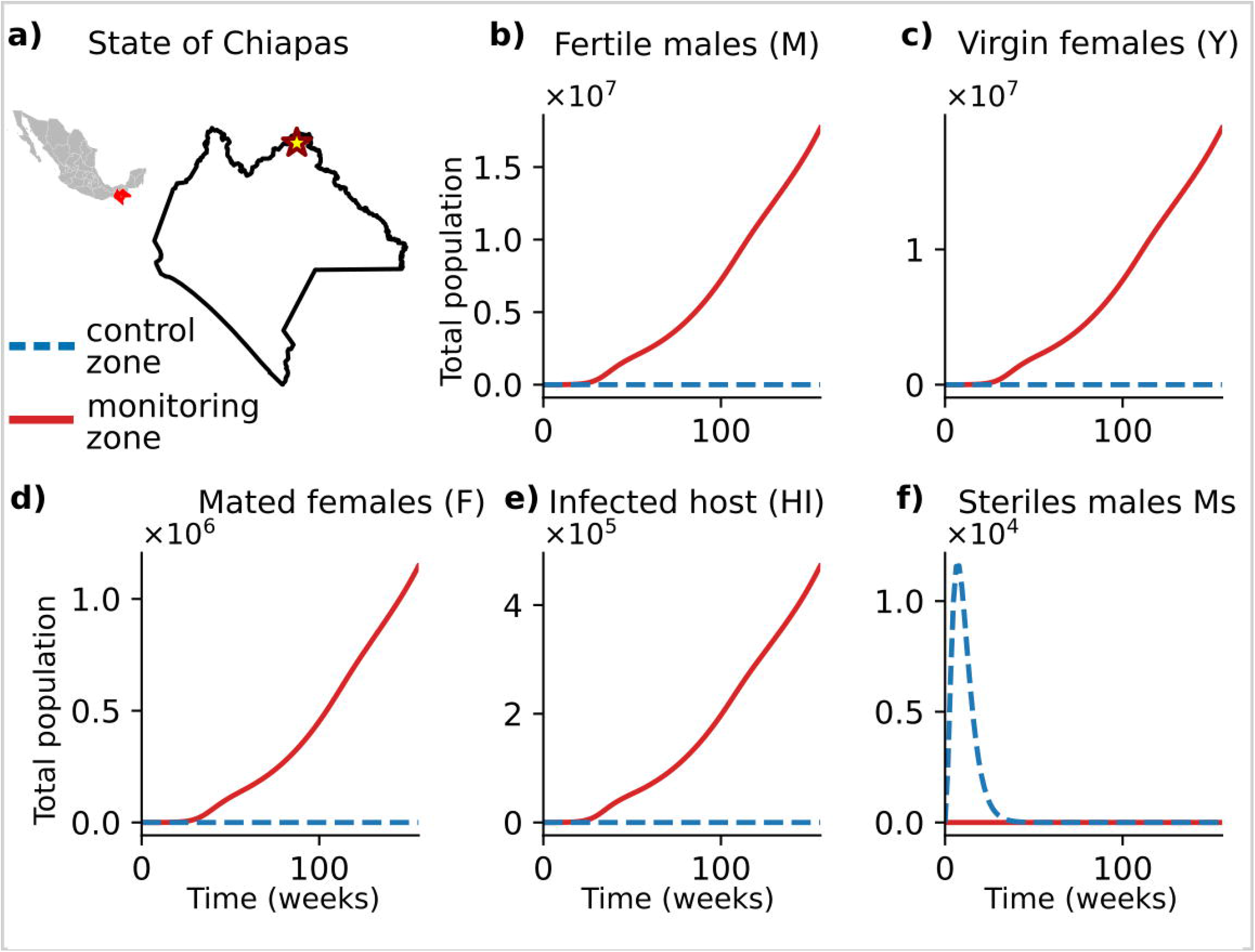
Comparison between observed data and model simulations. Comparison of the data reported by SENASICA and the results generated by our model using the optimized parameter set. Panel (a) shows the number of infected municipalities predicted by the model, while panel (b) shows the proportion of infected area relative to the total area of the state. In both panels, the dashed orange line represents the ensemble average of 50 independent simulations, the thin light-orange lines show individual model realizations tracking the variability introduced by stochastic dispersal events, and the shaded orange band denotes one standard deviation from the mean. Panel (c) compares the municipalities reported as infected by SENASICA in January 2026 with those simulated by our model. An exact replication of the spatial infection pattern is not expected, since the data are reported by municipalities regardless of the spatial extention of them. Also, the data seem to be incomplete, while our calculations consider long-distance infection events as stochastic. Nevertheless, the model is able to reproduce the overall spatial extent and distribution of the outbreak. Geographic boundaries were obtained from INEGI shapefiles and used for the spatial visualization after Python-based processing. These geographic data are publicly available and freely downloadable from INEGI.

The results show that in just over one year the infestation would have spread across the entire state, which is consistent with the current situation.

**S1 Video. Simulation of an initial infection event in Catazajá, Chiapas (yellow star)**. SENASICA surveillance data (left) are compared with the predictions obtained from our model (right). The model is solved on a 50 × 50 spatial grid. To obtain municipal-level results, the grid is intersected with municipal boundaries, and the values of all grid cells contained within each municipality are aggregated. Municipalities with a total number of infected animals *≤*1 are shown in red. Geographic boundaries were obtained from INEGI shapefiles and used for the spatial visualization after Python-based processing. These geographic data are publicly available and freely downloadable from INEGI.

### Evaluation of multiple scenarios

We evaluated the effectiveness of the control function in the spatial domain. When applied to every cell in the grid, it is possible to eradicate the infestation within the time estimated by the control function and to keep it suppressed even in the presence of re-outbreaks. These stochastic re-outbreaks are explicitly modeled to have the exact same population scale and magnitude as the initial seed cases. This indicates that the function, although optimized for a local model without spatial fluxes, remains effective when extended to the spatial case (Fig 6, dashed blue plots). In the absence of further-outbreaks, the extinction time is 46 weeks. When re-outbreaks are present, the pest remains controlled at minimal population levels. However, due to the continuous and distributed action of the feedback control, these new introductions are suppressed almost instantaneously, failing to exhibit any significant growth and remaining restricted to minimal levels. This indicates that the function, although optimized for a local model without spatial fluxes, remains highly effective and robust when extended to the spatial case (Fig 6, dashed blue plots). In the absence of further outbreaks, the extinction time is 46 weeks. When re-outbreaks are present, the pest population is successfully maintained at near-zero levels.

**Fig 6.**
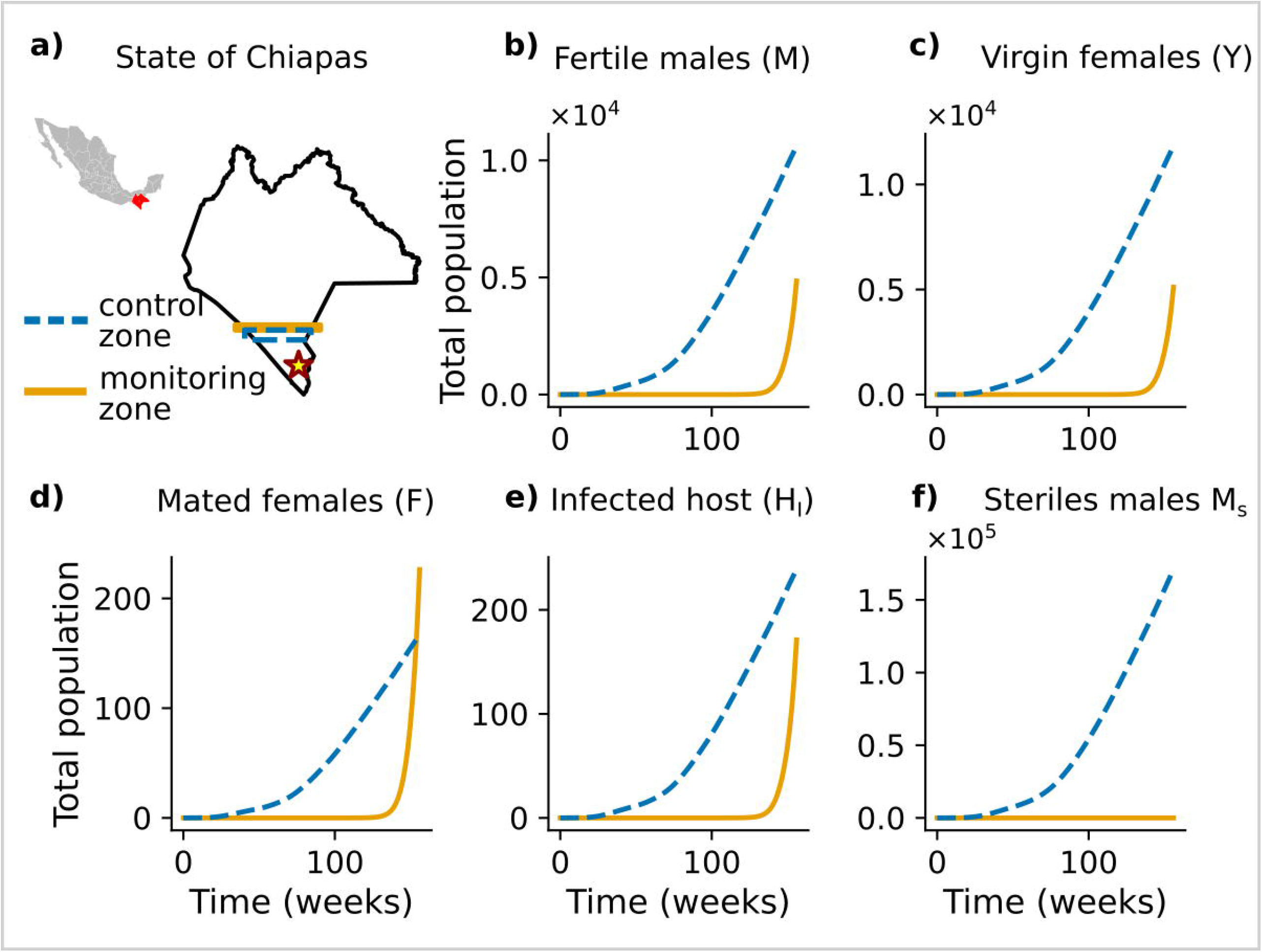
Pest dynamics with initial focus in Catazajá, northern Chiapas. (a) Geographic location of Catazajá, indicated by a yellow star. The inset map in the upper-left corner shows Mexico, with the state of Chiapas highlighted in red in the southern region bordering Guatemala. Dynamics of fertile males (b), virgin females (c), mated females (d), infected animals (e), and sterile males (f), for the uncontrolled case (red line) and the controlled case using our function for *u* (dashed blue line). Geographic boundaries were obtained from INEGI shapefiles and used for the spatial visualization after Python-based processing. These geographic data are publicly available and freely downloadable from INEGI.

We also analyzed the case in which the initial outbreak occurs in the southern part of the state, specifically in Tapachula, on the border with Guatemala, and containment is attempted through a control band approximately 24 km wide, where sterile males are released according to the control function. The results show that the infestation manages to penetrate the band in 3–5 weeks and, once crossed, spreads throughout the entire territory (see Fig 7)

**Fig 7.**
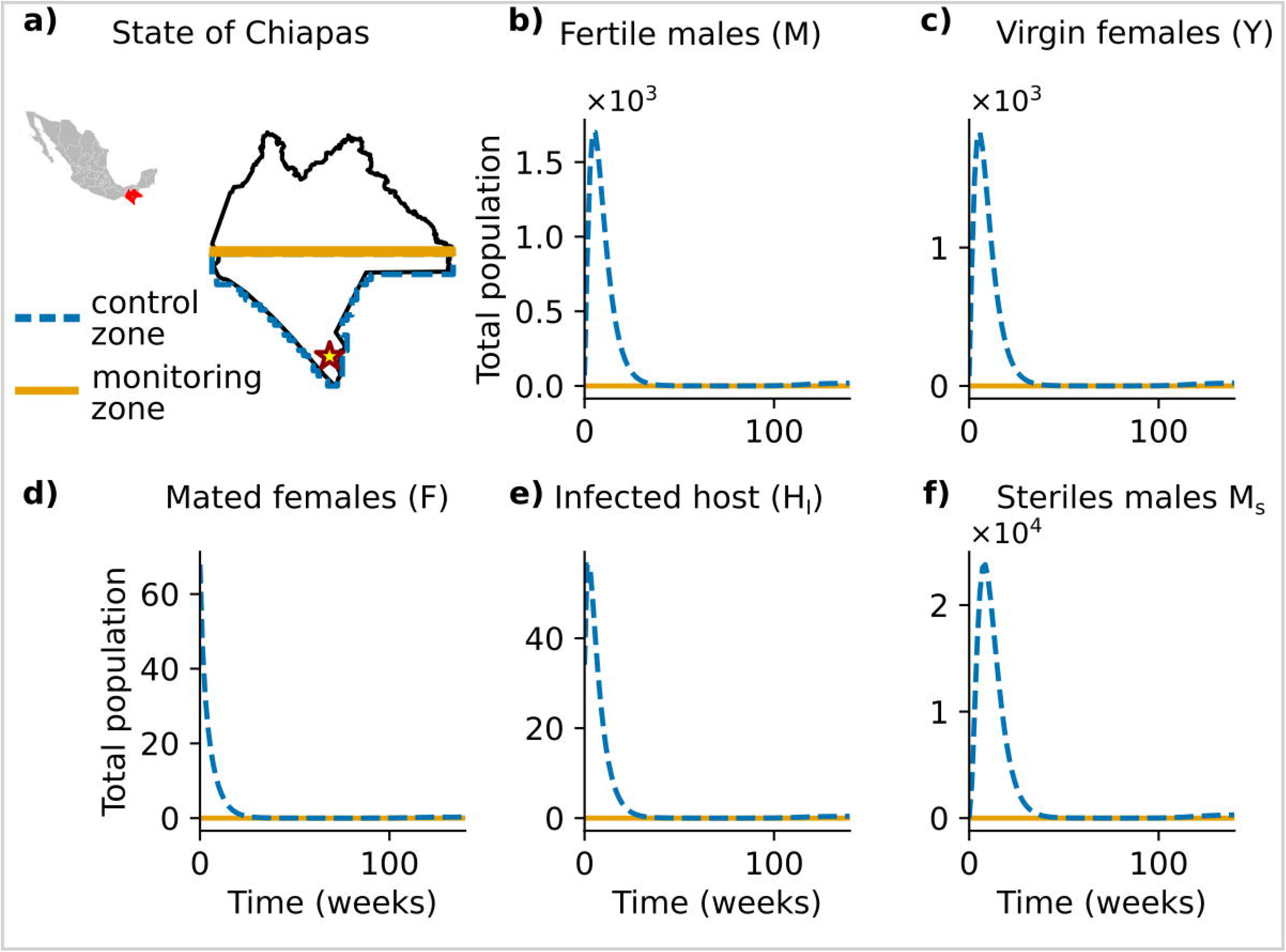
Simulation with an outbreak in Tapachula, near the Guatemala border. (a) Geographic location of Tapachula (yellow star). Sterile males are released only within the blue dotted rectangle (control zone), located immediately beyond the Tapachula border and approximately 24 km wide. The orange rectangle represents the monitoring zone, defined as the adjacent region where the pest population is monitored to determine whether the infestation crosses the control-zone boundary. Panels (b)–(f) show the dynamics of the total population summed over all pixels within the corresponding rectangles for fertile males (b), virgin females (c), mated females (d), infected hosts (e), and sterile males (f). The pest penetrates the control zone within 3–5 weeks and reaches the monitoring zone after approximately 10 weeks. Geographic boundaries were obtained from INEGI shapefiles and used for the spatial visualization after Python-based processing. These geographic data are publicly available and freely downloadable from INEGI.

Finally, we evaluated a scenario in which the infestation can be suppressed by releasing sterile males in the outbreak zone and within a radius of approximately 120 km around it, using the control function (Fig 8). Under these conditions, extinction is achieved in 47 weeks, consistent with the values reported in Table 3.

**Fig 8.**
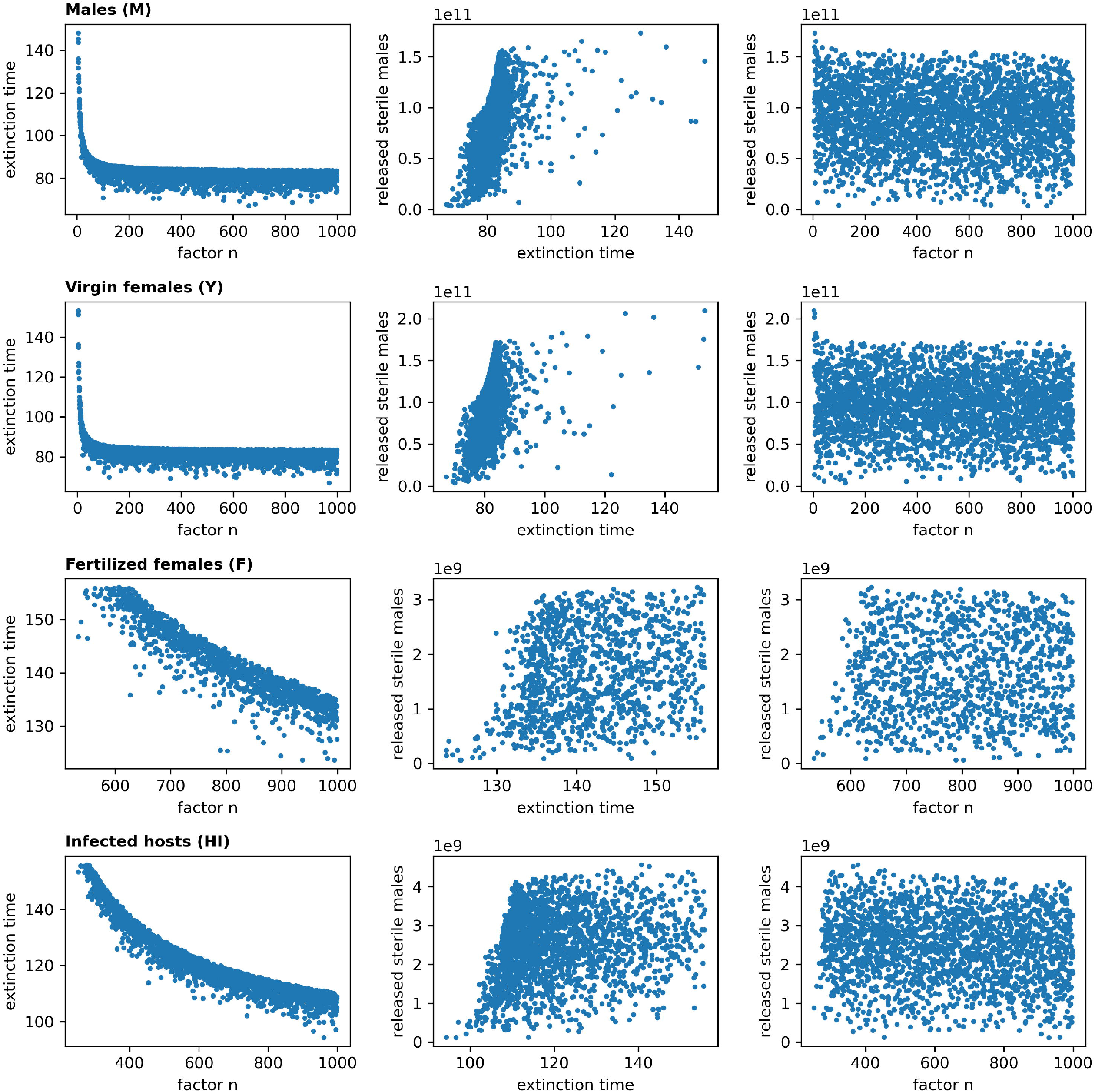
Eradication with sterile male releases at the focus and within a 120 km radius. (a) The outbreak is assumed to originate in Tapachula (yellow star), with sterile male releases applied within the focus area and in a region located approximately 120 km north of Tapachula (blue dashed polygon). The monitoring zone, where the pest population is evaluated to determine whether it crosses the control-zone boundary, is indicated by the orange rectangle. Under this strategy, the pest is eradicated from the control zone in approximately 47 weeks and never reaches the monitoring zone, as illustrated by the dynamics of fertile males (b), virgin females (c), mated females (d), infected animals (e), and sterile males (f). Geographic boundaries were obtained from INEGI shapefiles and used for the spatial visualization after Python-based processing. These geographic data are publicly available and freely downloadable from INEGI.

## Discussion

In this work, we developed a mathematical model tailored to describe the biology of the New World screwworm and proposed a feedback control function that determines the number of sterile males to release in order to suppress the infestation. This function adapts to the current infestation level and allows the exploration of trade-offs between the time required to eradicate the pest and the total number of sterile males deployed. Since direct field counts of wild males and virgin females are not feasible, we constructed an observer model that estimates these variables from the number of infected animals, consistent with the census methodology currently employed by SENASICA, and showed that, when coupled with the population dynamics model, it is sufficient to drive the pest to extinction.

We further extended the control function to a spatially explicit model incorporating dispersal between neighboring regions, and demonstrated that it retains its effectiveness, including comparable eradication timescales. Using the spatial model, we evaluated a containment strategy based on a sterile male release barrier and found that the pest can breach such a barrier within a few weeks. Nevertheless, eradication is achievable if the control effort is applied at the source of infestation and within a radius of approximately 120 km.

Unlike the number of sterile males required, which scales with the size of the infestation, the eradication time consistently falls within a range of 60 to 100 weeks. The analytical results from the simplified model formally confirm the observation that the local decay rates near the extinction equilibrium are governed exclusively by the pest’s mortality parameters *β*_*F*_ and *β*_*M*_, both of which are intrinsic biological constants. This result is consistent with the bifurcation structure of the full model in the absence of control, the system possesses three equilibria, extinction and carrying capacity (both stable), and an intermediate unstable threshold. Because this unstable threshold lies at a very low population level, any non-trivial infestation will grow toward carrying capacity. The release of sterile males shifts this threshold upward; once it exceeds the current infestation level, the system is attracted toward the extinction equilibrium.

A particularly notable outcome of the model calibration concerns the spatial spread dynamics. The calibration procedure identified an optimal diffusion coefficient of *D* = 0.002 (in grid-cell units per week), corresponding to a physical root-mean-square dispersal distance of approximately 0.72 km per week. This value is substantially lower than the empirical mark-recapture estimates reported by [18], which place the 90th-percentile dispersal radius between 1.25 and 2.85 km per observation period under favorable conditions, and up to 22 km under less favorable ones. The fact that the calibrated diffusion coefficient falls well below the biologically plausible range suggest that continuous Fickian diffusion is not the primary driver of spatial spread in this outbreak. Instead, the dominant mechanism appears to be the stochastic long-distance seeding term, which was essential to reproduce the observed spatio-temporal pattern of infected municipalities. This interpretation is consistent with field observations: under favorable environmental conditions *Cochliomyia hominivorax* females tend to remain near their oviposition sites and do not disperse widely by flight [18]. The observed jump-dispersal pattern is therefore more likely driven by human-mediated long-distance transport, in particular the movement of infested cattle and livestock products along inland road networks. This mechanistic interpretation aligns with the uniform spatial seeding assumption adopted in the model and underscores the importance of biosecurity measures targeting livestock transport as a complement to sterile male releases.

The present work shares its general motivation with recent studies on SIT-based feedback control for mosquito populations [10, 11], which demonstrated global stabilization of SIT models via Lyapunov functions. However, those formulations do not address the minimization of the total number of sterile insects released, which is a critical constraint when production capacity is limited. The present work focuses instead on numerically characterizing feedback gain matrices that achieve a favorable trade-off between the total number of sterile males deployed and the eradication time, providing explicit gain values for a range of infestation scenarios. This comes at the cost of local rather than global stability guarantees, since our analysis rests on linearizations of the nonlinear system at sampled points in the state space. Extending the framework to incorporate rigorous global stability proofs alongside resource constraints remains part of ongoing work.

Although the model parameters have been calibrated to the known biology of the screwworm, estimating the diffusion coefficient at a national scale remains challenging. Future extensions of the model could incorporate cattle mobility networks explicitly and the spatial heterogeneity of ecological niches that favor pest establishment and growth. The infestation has now spread across a large portion of Mexican territory, making it necessary to implement the model at a national rather than a Chiapas-specific scale. A critical open question is whether the total sterile male production capacity currently available in Mexico is sufficient to address the ongoing outbreak and, if so, what the Pareto-efficient spatial allocation strategy would be across the national territory. All these questions are the subject of ongoing research.

## Supporting Information

This document presents the explicit steady-state solution of the mathematical model described by Eqs. (1)–(5).

## Acknowledgments

R.R. acknowledges the Instituto Politécnico Nacional for the projects 20253592-20260826 and the Secretaría de Ciencia, Humanidades, Tecnología e Innovación for the postdoctoral fellowship at the Instituto de Física, UNAM.

Both authors are grateful to Francisco Infante, José Luis Quintero Fong, Luis A. Cisneros-Ake, Vincenzo Bertolini and Raúl Vera for fruitful discussions.

## Notes

### Competing Interest Statement

The authors have declared no competing interest.

