## Supporting information for "A mathematical model for the efficient control of the New World screwworm"

### 1 Equilibria for the complete model

The steady-state equations of the model are

$$(1-r)\mu H_I - \beta_M M + \Lambda_M = 0 \tag{1}$$

$$r\mu H_I - \gamma_1 Y\frac{M}{M+M_s} - \gamma_2 Y\frac{M_s}{M+M_s} - \beta_Y Y + \Lambda_Y = 0 \tag{2}$$

$$\gamma_1 Y\frac{M}{M+M_s} - \beta_F F + \Lambda_F = 0 \tag{3}$$

$$\alpha_I F\left(1-\frac{H_I}{K_I}\right) - \beta_I H_I = 0 \tag{4}$$

$$u - \beta_s M_s = 0 \tag{5}$$

and we solve the system and obtain the following solutions.

We obtain three equilibria. The first is the trivial equilibrium:

$$\begin{pmatrix} 0 & 0 & 0 & 0 & \frac{5u}{2} \end{pmatrix} \tag{6}$$

The second:

$$\begin{pmatrix} \frac{12679\,K_I}{880} - \frac{\sqrt{K_I\,(160757041\,K_I - 18663700\,u)}}{880} \\ \frac{480500\,K_I\left(\frac{409\,K_I}{1100} - \frac{\sqrt{K_I\,(160757041\,K_I-18663700\,u)}}{34100}\right) + 235\,K_I\,u + 31000\,u\left(\frac{409\,K_I}{1100} - \frac{\sqrt{K_I\,(160757041\,K_I-18663700\,u)}}{34100}\right)}{11532\,K_I + 722\,u} \\ \frac{1057100\,K_I\left(\frac{409\,K_I}{1100} - \frac{\sqrt{K_I\,(160757041\,K_I-18663700\,u)}}{34100}\right) - 16967\,K_I\,u}{403620\,K_I + 25270\,u} \\ \frac{409\,K_I}{1100} - \frac{\sqrt{K_I\,(160757041\,K_I-18663700\,u)}}{34100} \\ \frac{5u}{2} \end{pmatrix} \tag{7}$$

The third:

$$\left(\begin{array}{c} \frac{12679\,K_I}{880}+\frac{\sqrt{K_I\,\left(160757041\,K_I-18663700\,u\right)}}{880}\\ \frac{480500\,K_I\,\left(\frac{409\,K_I}{1100}+\frac{\sqrt{K_I\,\left(160757041\,K_I-18663700\,u\right)}}{34100}\right)+235\,K_I\,u+31000\,u\,\left(\frac{409\,K_I}{1100}+\frac{\sqrt{K_I\,\left(160757041\,K_I-18663700\,u\right)}}{34100}\right)}{11532\,K_I+722\,u}\\ \frac{1057100\,K_I\,\left(\frac{409\,K_I}{1100}+\frac{\sqrt{K_I\,\left(160757041\,K_I-18663700\,u\right)}}{34100}\right)-16967\,K_I\,u}{403620\,K_I+25270\,u}\\ \frac{409\,K_I}{1100}+\frac{\sqrt{K_I\,\left(160757041\,K_I-18663700\,u\right)}}{34100}\\ \frac{5\,u}{2} \end{array}\right) \tag{8}$$

In the second and third equilibria, the discriminant $\Delta = K_I(160{,}757{,}041\,K_I - 18{,}663{,}700\,u)$ appears in the equilibrium expressions.
